# Preclinical safety assessment of Koomba kaat 1 and Biyabeda mokiny 1 phages for respiratory application against *Staphylococcus aureus*

**DOI:** 10.64898/2026.07.24.740517

**Authors:** Joshua J Iszatt, Alexander N Larcombe, Luke W Garratt, Stephen M Stick, Anthony Kicic, WAERP and Phage WA

## Abstract

Therapeutic bacteriophages are promising alternatives to antibiotics for treating antimicrobial-resistant bacterial infections. For pulmonary infections, direct respiratory delivery offers therapeutic advantages; however, preclinical evaluation of respiratory safety remains underdeveloped. Bacteriophages exhibit genomic and biological diversity, and safety cannot be assumed for individual candidates. Here, we evaluated the respiratory safety of lytic *Staphylococcus aureus* bacteriophages, Koomba kaat 1 and Biyabeda mokiny 1, using human and murine preclinical models.

Differentiated primary airway epithelial cells derived from six healthy paediatric donors were exposed apically to purified phage (1 × 10^9^ PFU/mL) for 24 hours. Barrier integrity, epithelial morphology, mucus production, cytotoxicity, and interleukin-8 release were assessed. Safety was further investigated in adult C57BL/6J mice receiving intranasal phage administration (1 × 10^9^ PFU) twice daily (14 days).

Clinical observations, body weight, organ pathology, blood biochemistry, bronchoalveolar lavage cellularity, protein concentration, and inflammatory mediators were evaluated. Neither phage altered epithelial morphology, barrier integrity, mucus production, cytotoxicity, nor inflammatory responses in differentiated cultures. Repeated intranasal administration was well tolerated in vivo, with no adverse clinical signs, weight loss, macroscopic pathology, or treatment-associated changes in pulmonary cellularity or tissue histopathology. Differences in blood biochemistry and inflammatory mediators were small and not accompanied by epithelial injury or pulmonary inflammation.

Collectively, these findings demonstrate that Koomba kaat 1 and Biyabeda mokiny 1 exhibit favourable safety profiles following repeated airway administration. This study establishes a comprehensive framework for the preclinical respiratory safety assessment of bacteriophages and provides support for the clinical development of inhaled phage for *S. aureus* respiratory infections.

## Introduction

Antimicrobial resistance (AMR) is one of the greatest threats to global health, with lower respiratory tract infections (LRTIs) remaining a leading cause of AMR-associated morbidity and mortality (Naghavi et al., 2024). *Staphylococcus aureus* (*S. aureus*) is an important respiratory pathogen responsible for both community- and healthcare-associated pneumonia and contributes substantially to chronic airway infection in vulnerable populations, including individuals with cystic fibrosis (Chambers & Deleo, 2009; Heltshe et al., 2015). Specifically, *S. aureus* is also a significant cause of LRTIs, including pneumonia, particularly in vulnerable populations and in both hospital- and community-acquired settings (Tong et al., 2015). The emergence of antibiotic-resistant strains including methicillin-resistant *S. aureus* (MRSA) within hospital and community settings has led to increasing efforts to find non-antibiotic treatment alternatives (Czaplewski et al., 2016).

Lytic bacteriophages (phages) have re-emerged as promising therapeutics because of their ability to selectively infect and kill bacterial pathogens (Cui et al., 2017; A. C. C. de Melo et al., 2019; Estrella et al., 2016). Clinical experience has demonstrated their potential as stand-alone treatments, defined phage cocktails, or adjuncts to antibiotics, where synergistic activity may improve bacterial clearance and reduce the emergence of antimicrobial resistance (Aslam et al., 2020; Gainey et al., 2020; Law et al., 2019; Molina et al., 2021, 2022). For respiratory infections, direct airway delivery via inhalation or intranasal administration offers the advantage of achieving high local phage concentrations while limiting systemic exposure (Iszatt, 2023). Despite increasing clinical interest, respiratory-specific safety assessment remains an underdeveloped aspect of phage therapeutic translation. Comprehensive reviews have highlighted that relatively few studies have systematically investigated pulmonary toxicity, epithelial responses, or inflammatory effects following respiratory administration of purified phages (Liu et al., 2021). Unlike conventional therapeutics, phages exhibit substantial genomic and biological diversity, meaning that safety cannot be inferred across related viruses or predicted solely by genomic analyses (Cook et al., 2021; Danis-Wlodarczyk et al., 2016; Dion et al., 2020; Lehman et al., 2019; L. D. R. Melo et al., 2020). Consequently, each therapeutic phage requires empirical safety evaluation before clinical application.

Current preclinical studies are further limited by the widespread use of immortalised airway cell lines cultured as submerged monolayers, models that poorly represent the differentiated airway epithelium, epithelial polarity, mucus production, and barrier integrity observed *in vivo* (Martinovich et al., 2017; Pezzulo et al., 2011; Silva et al., 2023; Trend et al., 2018). Similarly, relatively few *in vivo* studies have comprehensively evaluated repeated respiratory administration of purified phages in the absence of bacterial infection while assessing systemic physiology, pulmonary inflammation, and tissue pathology (Skronska-Wasek et al., 2021).

We previously isolated and characterised two lytic *S. aureus* bacteriophages, Koomba kaat 1 (Silviavirus) and Biyabeda mokiny 1 (Kayvirus), demonstrating favourable host range, biofilm disruption, stability, and the absence of recognised lysogeny-associated genes, antimicrobial resistance determinants, and bacterial virulence factors (Iszatt et al., 2025). Here, we evaluated their intrinsic respiratory safety using complementary human and murine preclinical models. Differentiated primary airway epithelial cultures were used to assess epithelial integrity, morphology, mucus production, cytotoxicity, and inflammatory responses, while a repeated-dose murine model examined clinical tolerance, pulmonary inflammation, systemic physiology, and tissue pathology following twice-daily intranasal administration for 14 days, reflecting clinically relevant treatment durations (Petrovic Fabijan et al., 2020). We hypothesised that both phages would be well tolerated following respiratory administration, providing critical preclinical evidence to support their clinical translation and contributing towards a standardised framework for respiratory safety assessment of inhaled phage therapeutics.

## Materials and Methods

### Phage preparation and characterisation

The lytic *Staphylococcus aureus* bacteriophages Koomba kaat 1 and Biyabeda mokiny 1 were propagated, purified, and prepared from a single production batch as previously described (Iszatt et al., 2025). Clinical respiratory MRSA isolates SA09 and SA01 (Provided by Professor Scott Bell, the Queensland Institute of Medical Research, QIMR Berghofer, Queensland, Australia) served as propagation hosts for Koomba kaat 1 and Biyabeda mokiny 1, respectively. Bacteria were cultured overnight in Tryptic Soy (TS) broth (BD Difco™) at 37°C with agitation (120 rpm) and maintained as glycerol stocks (25% v/v) at -80°C. High-titre lysates were filter sterilised (0.22 μm) before purification by high-performance liquid chromatography using a HiTrap® BIA Monolithic Column (BIA Separations) on an ÄKTA pure™ chromatography system (Cytiva). Fractions corresponding to the principal UV280 peaks were collected, pooled and used for all experiments. The absence of *S. aureus* enterotoxins A–E from purified phage preparations was confirmed using the RIDASCREEN® enzyme immunoassay (R-Biopharm) according to the manufacturer’s instructions (limit of detection 0.25 ng/mL), all samples used tin this study were below the limit of detection. Heat-killed bacterial suspensions were prepared as inflammatory comparators. Mid-logarithmic-phase cultures (∼1 × 10^9^ CFU/mL) were washed three times with sterile phosphate-buffered saline (PBS), heat inactivated (80°C, 60 min) and confirmed non-viable by overnight culture on TS agar and in TS broth.

### Primary airway epithelial cell model

Primary airway epithelial cells (pAECs) were obtained from six healthy children (three males, three females) enrolled in the Western Australian Epithelial Research Program (WAERP; ethics approval #901). Cells were expanded and differentiated at air-liquid interface (ALI) on Transwell® inserts for 28 days as previously described (Martinovich et al., 2017). Differentiation was confirmed by transepithelial electrical resistance (TEER), with cultures demonstrating a mean resistance of 740 ± 111 Ω/cm^2^ prior to experimentation. Differentiated cultures received a single apical application (10 μL) of PBS, Koomba kaat 1 (1 × 10^9^ PFU/mL), Biyabeda mokiny 1 (1 × 10^9^ PFU/mL), heat-killed SA01 (1 × 10^8^ CFU/mL), or heat-killed SA09 (1 × 10^8^ CFU/mL). Following 24 hours of incubation, apical washings and basolateral media were collected for downstream analyses. TEER measurements were repeated immediately before exposure and after 24 hours to assess epithelial barrier integrity. Epithelial inflammation was quantified by measuring interleukin-8 (IL-8) using ELISA (BD Biosciences) according to previously described methods (Kicic et al., 2006). IL-8 concentrations were normalised to total cellular protein determined by bicinchoninic acid (BCA) assay (Thermo Fisher Scientific). Cytotoxicity was assessed by lactate dehydrogenase (LDH) release using the CytoTox 96® Non-Radioactive Cytotoxicity Assay (Promega). To assess epithelial morphology and mucus production, inserts were fixed in 10% neutral buffered formalin, paraffin embedded, sectioned (5 μm), and stained with haematoxylin and eosin or Alcian blue. Mucus production was quantified using BIMANA software as the proportion of Alcian blue-positive pixels relative to total epithelial tissue area (Alphons Gwatimba, 2023).

### Murine respiratory safety model

Thirty-six adult C57BL/6J mice (18 males, 18 females; ∼8 weeks old) were obtained from the Animal Resources Centre (Murdoch, Australia) and housed under standard conditions with unrestricted access to food and water. All procedures were approved by The Kids Research Institute Australia Animal Ethics Committee (P2267). Mice received intranasal administration of PBS, Koomba kaat 1, or Biyabeda mokiny 1 (1 × 10^9^ PFU in 50 μL PBS) twice daily for 14 consecutive days under isoflurane anaesthesia. Six males and six females were included in each treatment group. Body weight, food and water consumption, and clinical wellbeing were monitored throughout the study. Approximately 20 hours after the final treatment, mice were anaesthetised with ketamine/xylazine, mechanically ventilated, and blood collected by cardiac puncture for blood gas and biochemical analyses (i-STAT Alinity, Abbott Laboratories). A complete necropsy was performed, with visual examination of major organs and measurement of liver, spleen and kidney weights. Bronchoalveolar lavage fluid (BALF) was collected by three sequential saline washes (0.5 mL each). Total and differential cell counts were determined following cytospin preparation and Rapid Stain staining. BALF total protein was measured using a Pierce™ BCA Protein Assay (Thermo Fisher Scientific). Inflammatory mediators were quantified using the Bio-Plex Pro™ Mouse Cytokine 23-Plex assay (Bio-Rad) following the manufacturer’s recommendations. BALF processing was performed as previously described (Landwehr et al., 2023).

### Statistical analysis

All *in vitro* experiments were performed using primary airway epithelial cells from 6 biological donors (3 male, 3 female). *In vivo* experiments used 6 mice per sex per treatment group. Statistical analyses were conducted using GraphPad Prism v8.4.3 (GraphPad Software, La Jolla, CA, USA). Comparisons containing mixed model analyses were performed using 2-way ANOVA. P-values <0.05 were considered statistically significant for all tests unless specified otherwise. Mucus production and apical and basolateral LDH and IL-8 fold changes were assessed by one-way ANOVA; LDH treatments were compared to the PBS baseline by one-sample t-test and IL-8 phages to their host strain using Šídák-corrected comparisons. TEER was assessed by Kruskal-Wallis with Dwass-Steel-Critchlow-Fligner post-hoc comparisons. Body weight was compared between Day 1 and Day 15 within each sex by paired t-test.

Differential cell counts, blood gases and chemistry, whole protein quantitation, and organ weight data (spleen, liver, kidneys) were all assessed using two-way ANOVA with sex and treatment as factors. Data were transformed where required to satisfy the assumptions of normality and homogeneity of variance. Protein data were filtered for outliers that were > 2x standard deviations away from the group mean. Statistical tests were run using Jamovi (2023) version 2.3 (2022 release). Bioplex raw data were filtered to remove individual mediators where every value fell below the detection limit and remaining data were processed by removing individual outlier measurements that lay more than 2 standard deviations outside of the group mean. Remaining data points that fell below the detection range were changed to half of the lowest standard to enable subsequent statistical analysis as previously described (Landwehr et al., 2023). To assess differences between males, females, phage treatments, and controls, a Generalised Linear Model (GLM) approach was performed with the Python modules statsmodels (v0.13.4) and SciPy v1.11.0 (Virtanen et al., 2020).

## Results

### Koomba kaat 1 and Biyabeda mokiny 1 do not alter epithelial morphology, barrier integrity or mucus production

Differentiated primary airway epithelial cell (pAEC) cultures remained morphologically intact following 24-hour apical exposure to Koomba kaat 1, Biyabeda mokiny 1, heat-killed *S. aureus* (SA01 or SA09), or PBS control (Figure 1). Histological examination revealed a well-differentiated pseudostratified epithelium with no evidence of epithelial disruption, cellular detachment, or morphological abnormalities in any treatment group. Similarly, Alcian blue staining demonstrated no detectable effect of phage exposure on epithelial mucus production (Figure 2). Quantitative image analysis identified no overall difference between treatment groups (one-way ANOVA, *P* = 0.535). Relative to PBS controls (0.092 ± 0.062 stained pixels/tissue area), neither Koomba kaat 1 (0.043 ± 0.025; *P* = 0.715) nor Biyabeda mokiny 1 (0.088 ± 0.087; *P* = 1.000) altered mucus production. Heat-killed bacterial controls likewise did not differ from PBS (SA01: 0.106 ± 0.070, *P* = 0.996; SA09: 0.097 ± 0.073, *P* = 1.000). Epithelial barrier integrity, assessed by transepithelial electrical resistance (TEER), was maintained following phage exposure. No significant differences were detected between treatment groups (*P* = 0.751). TEER values following exposure to Koomba kaat 1 (519 ± 353 Ω/cm^2^) and Biyabeda mokiny 1 (384 ± 402 Ω/cm^2^) were comparable with PBS controls (409 ± 385 Ω/cm^2^), while heat-killed SA01 (352 ± 421 Ω/cm^2^) and SA09 (546 ± 394 Ω/cm^2^) similarly produced no measurable changes. Collectively, these findings indicate that neither therapeutic phage adversely affected epithelial morphology, mucus production, or barrier integrity following high-dose apical exposure.

**Figure 1:**
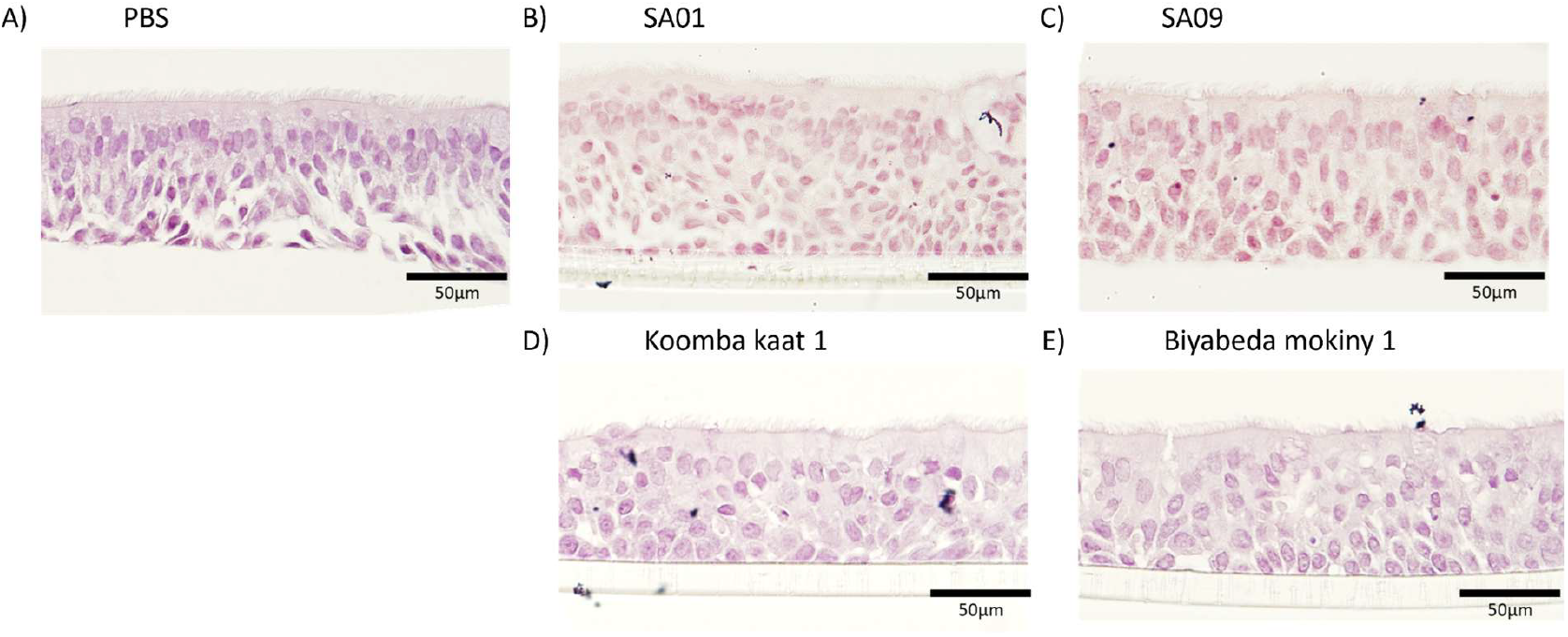
Phage administration did not alter airway epithelial structure. Primary airway epithelial cells from healthy children (n=6) were cultured at the air-liquid interface and exposed apically for 24 hours to PBS (A), Koomba kaat 1 (B), Biyabeda mokiny 1 (C), heat-inactivated SA01 (D), or heat-inactivated SA09 (E). Inserts were fixed in 10% neutral buffered formalin, sectioned (5 μm), and stained with haematoxylin and eosin. Images shown are representative, taken at 40× magnification. No visible differences in epithelial architecture were observed between phage- or bacteria-exposed cultures (B–E) and the PBS control (A).

**Figure 2:**
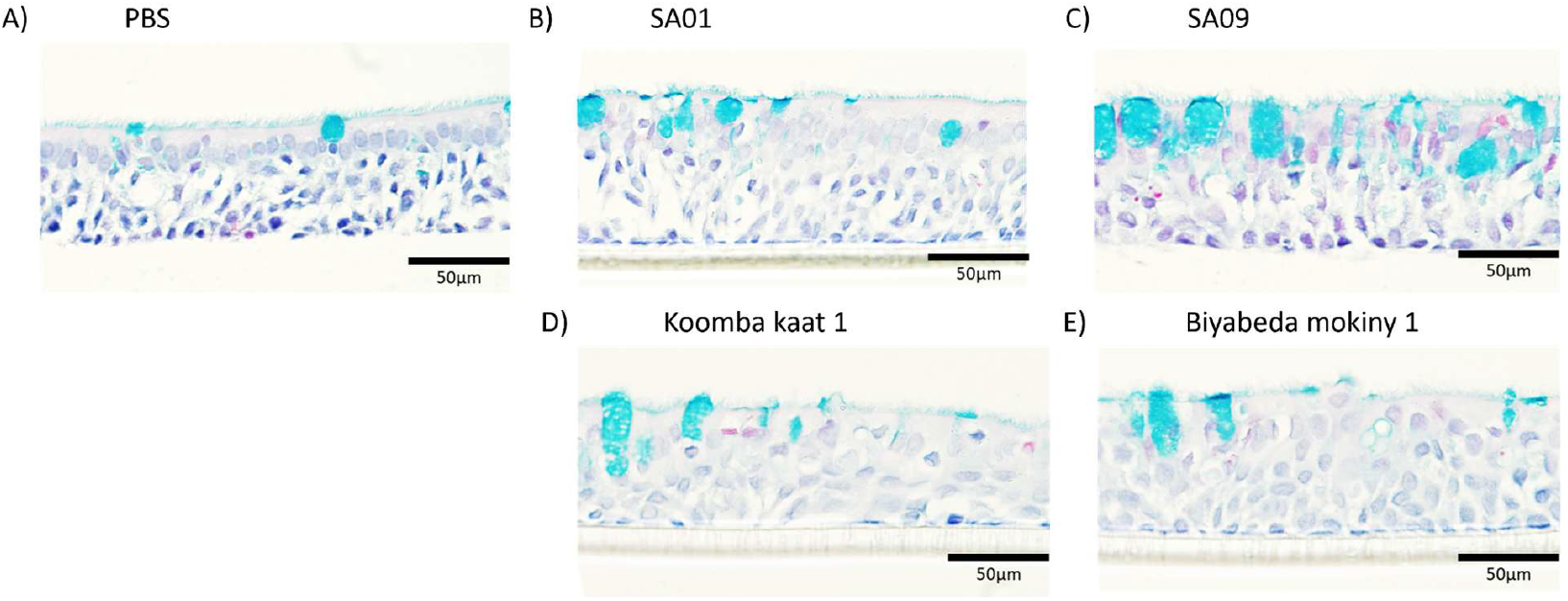
Phage administration did not induce mucus production. Primary airway epithelial cells from healthy children (n=6) were cultured at the air-liquid interface and exposed apically for 24 hours to PBS (A), Koomba kaat 1 (B), Biyabeda mokiny 1 (C), heat-inactivated SA01 (D), or heat-inactivated SA09 (E). Inserts were fixed in 10% neutral buffered formalin, sectioned (5 μm), and stained with Alcian blue. Mucus content was quantified using BIMANA as the ratio of Alcian blue-stained pixels to total tissue area. Images shown are representative, taken at 40x magnification.

### Therapeutic phages do not induce epithelial cytotoxicity or clinically meaningful inflammatory responses

To determine whether phage exposure induced epithelial injury, lactate dehydrogenase (LDH) release was quantified in both apical washings and basolateral media following 24-hour exposure (Figure 3A,B). No overall differences were observed between treatment groups in either compartment (apical: *P* = 0.230; basolateral: *P* = 0.651). Relative to PBS controls, neither Koomba kaat 1 (apical: 2.85 ± 2.75-fold, *P* = 0.160; basolateral: 1.23 ± 1.29-fold, *P* = 0.683) nor Biyabeda mokiny 1 (apical: 0.87 ± 0.73-fold, *P* = 0.672; basolateral: 0.66 ± 0.75-fold, *P* = 0.320) increased LDH release. Heat-killed bacterial preparations also produced no significant increase in cytotoxicity compared with PBS (SA01 apical: *P* = 0.222, basolateral: *P* = 0.917; SA09 apical: *P* = 0.094, basolateral: *P* = 0.407).

**Figure 3:**
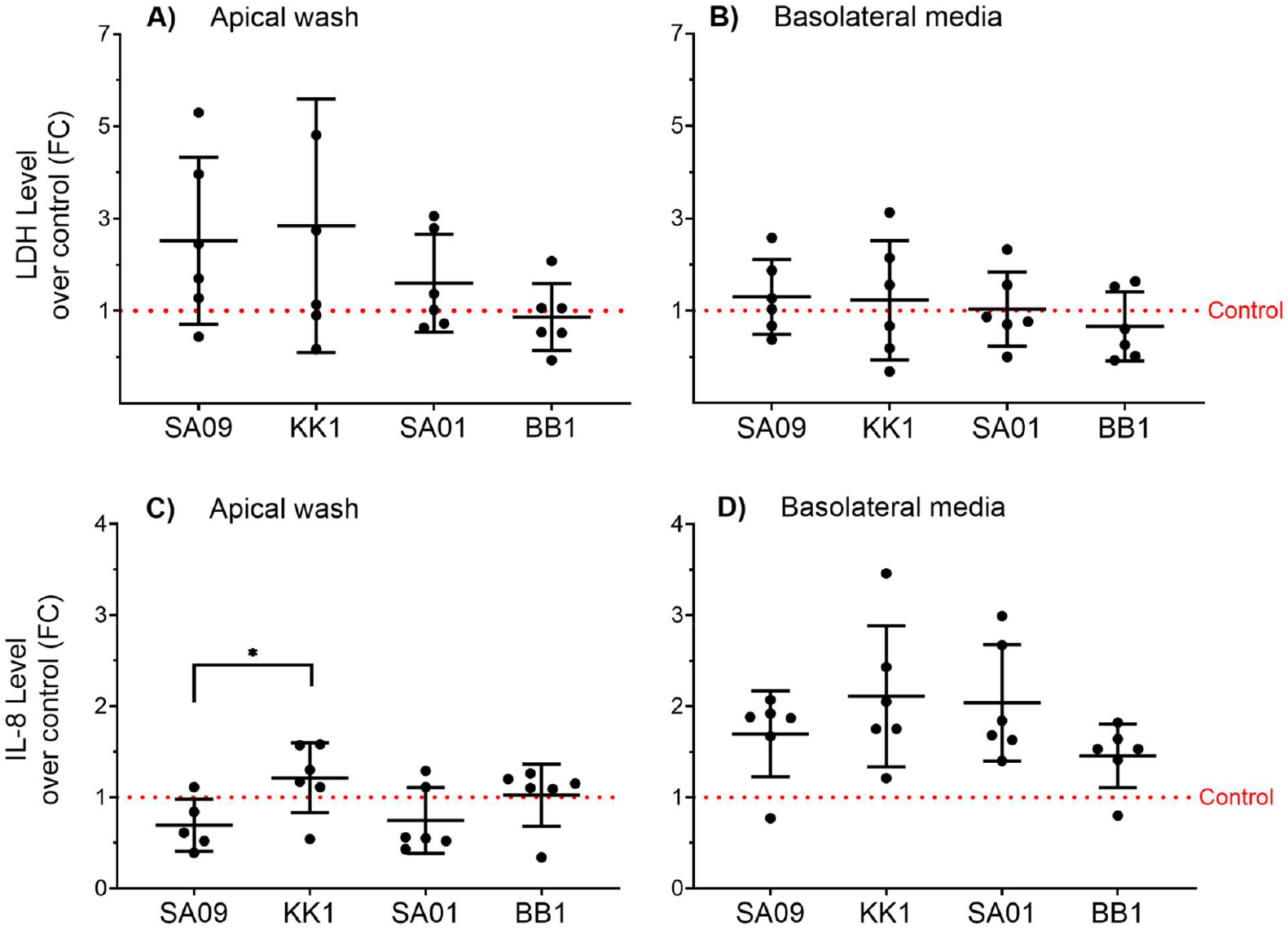
Koomba kaat 1 and Biyabeda mokiny 1 are not cytotoxic to the airway epithelium. Primary airway epithelial cells from healthy children (n=6) were cultured at the air-liquid interface and exposed apically for 24 hours to PBS, Koomba kaat 1 (KK1), Biyabeda mokiny 1 (BB1), heat-inactivated SA01, or heat-inactivated SA09. Apical washings and basolateral media were collected and assayed for LDH release (A, B) and IL-8 (C, D). IL-8 data were normalised to total protein determined by BCA assay. All data are presented as mean fold change relative to the PBS control (dashed line) ± standard deviation.

Inflammatory responses were assessed by measuring IL-8 secretion from the apical and basolateral compartments (Figure 3C,D). No overall treatment effect was detected for apical (*P* = 0.068) or basolateral (*P* = 0.212) IL-8 concentrations. Apical IL-8 responses remained low across all treatment groups, with fold changes of 1.21 ± 0.38 for Koomba kaat 1, 1.02 ± 0.34 for Biyabeda mokiny 1, 0.74 ± 0.36 for heat-killed SA01, and 0.69 ± 0.29 for heat-killed SA09 relative to PBS controls. Planned comparisons between each phage and its corresponding bacterial host identified a modest increase in apical IL-8 following Koomba kaat 1 exposure compared with heat-killed SA09 (mean difference 0.52; *P* = 0.047), whereas Biyabeda mokiny 1 did not differ from its propagation host SA01 (*P* = 0.326). Basolateral IL-8 concentrations were similarly unchanged, with comparable responses observed for Koomba kaat 1 (2.11 ± 0.77-fold), Biyabeda mokiny 1 (1.46 ± 0.35-fold), heat-killed SA01 (2.04 ± 0.64-fold), and heat-killed SA09 (1.70 ± 0.47-fold). Neither phage differed significantly from its corresponding bacterial comparator (Koomba kaat 1 versus SA09, *P* = 0.414; Biyabeda mokiny 1 versus SA01, *P* = 0.189). Overall, exposure of differentiated primary airway epithelial cultures to purified Koomba kaat 1 or Biyabeda mokiny 1 did not induce measurable epithelial injury or a biologically meaningful inflammatory response.

### Koomba kaat 1 and Biyabeda mokiny 1 are well tolerated following repeated respiratory administration

Repeated intranasal administration of Koomba kaat 1 or Biyabeda mokiny 1 twice daily for 14 days was well tolerated by C57BL/6J mice. No treatment-related adverse events or abnormalities in clinical observations were recorded throughout the study in either sex. Body weight remained stable during the treatment period (Figure 4). Neither Koomba kaat 1 nor Biyabeda mokiny 1 altered body weight relative to PBS-treated controls in male (*P* = 0.223) or female (*P* = 0.338) mice. Male mice demonstrated a modest physiological increase in body weight over the 14-day study (+0.6 g; paired *P* < 0.001), whereas female body weight remained unchanged (*P* = 0.194). Food and water consumption were comparable across all treatment groups (data not shown). Gross necropsy identified no abnormalities of the respiratory tract or other major organs, including the heart, liver, kidneys, spleen, gastrointestinal tract or reproductive organs. Likewise, phage administration had no effect on liver (*P* = 0.969), spleen (*P* = 0.561), left kidney (*P* = 0.372) or right kidney (*P* = 0.415) weights. As expected, males exhibited greater liver and kidney weights than females (both *P* < 0.001), reflecting normal sex-related physiological differences.

**Figure 4:**
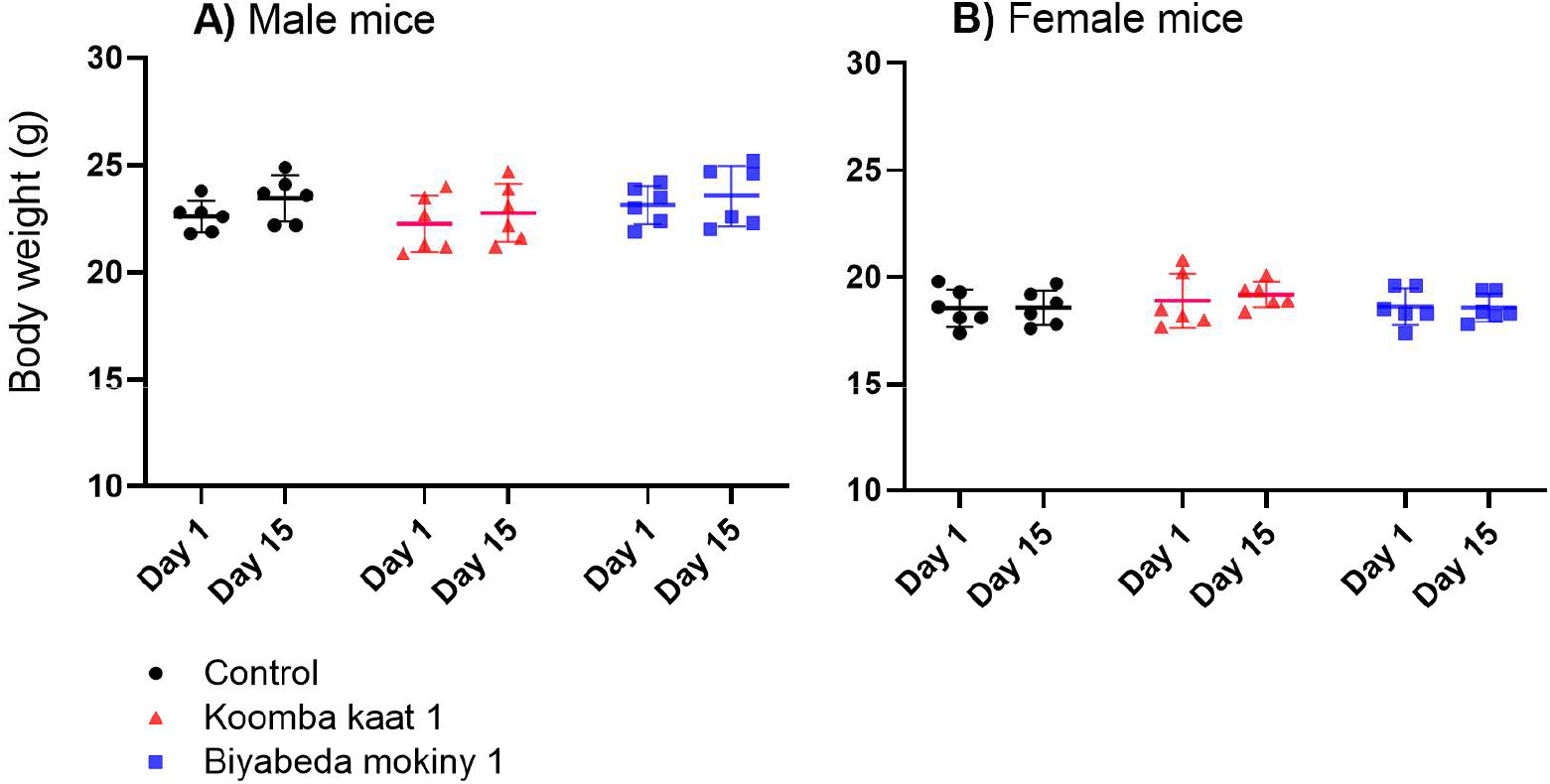
No reduction in body weight observed in mice after 14 days of twice a day phage treatment. Male (A) and female (B) mice weights on Day 1 and Day 15 for each treatment group. Male mice gained slightly over the 14 days, with no difference between treatment groups; female weights were unchanged.

### Blood biochemistry and pulmonary cellular responses remain unchanged following phage administration

Blood gas, biochemical and haematological analyses identified few treatment-associated differences (Supplementary Table 1). Blood glucose concentrations were lower in males receiving Biyabeda mokiny 1 than sex-matched PBS controls (17.98 ± 2.33 versus 22.37 ± 2.13 mmol/L; *P* = 0.038), whereas no differences were observed following Koomba kaat 1 administration. Base excess differed between sexes (males: −6.33 ± 2.63 mmol/L; females: −8.18 ± 2.63 mmol/L; *P* = 0.011), independent of treatment (*P* = 0.735). No additional blood gas, biochemical or haematological parameters differed between treatment groups.

Bronchoalveolar lavage fluid (BALF) analysis similarly demonstrated no evidence of treatment-associated cellular inflammation (Table 1). Total BALF cellularity, macrophage numbers and neutrophil counts did not differ between treatment groups (total cells: *P* = 0.473; macrophages: *P* = 0.520; neutrophils: *P* = 0.146), and no eosinophils or other inflammatory cell populations were detected. As no sex effects were identified for BALF cellularity or protein concentration, data were pooled for subsequent analyses.

**Table 1.** Total cells, macrophages, neutrophils, and total BALf protein for each treatment group.

| Treatment group | Total (cells/mL) | Macrophages (cells/mL) | Neutrophils (cells/mL) | BALf protein (µg/mL) |
| --- | --- | --- | --- | --- |
| Control | 30,097 ± 11,030 | 29,371 ± 10,905 | 727 ± 646 | 306 ± 49 |
| Koomba kaat 1 | 25,199 ± 10,186 | 24,795 ± 9,871 | 403 ± 403 | 313 ± 32 |
| Biyabeda mokiny 1 | 29,909 ± 9,093 | 29,144 ± 8,636 | 765 ± 981 | 247 ± 35 |

Total BALF protein differed between treatment groups (*P* = 0.002), with lower concentrations observed following Biyabeda mokiny 1 administration (247 ± 35 μg/mL) compared with PBS controls (306 ± 49 μg/mL) and Koomba kaat 1 (313 ± 32 μg/mL).

### Respiratory administration induces minimal changes in inflammatory mediator profiles

Of the 23 inflammatory mediators assessed in BALF, 18 were detected above the assay limit of quantification (Supplementary Table 2). Relative to sex-matched PBS controls, neither Koomba kaat 1 nor Biyabeda mokiny 1 altered concentrations of the principal pro-inflammatory mediators IL-1α, IL-6, G-CSF, KC or TNF-α. Biyabeda mokiny 1 did not significantly affect any of the 18 detectable mediators. In contrast, Koomba kaat 1 produced significant changes in three cytokines. IL-17 concentrations were reduced relative to controls (*P* = 0.017), with the reduction more pronounced in females (0.52 ± 0.24 pg/mL versus 0.83 ± 0.11 pg/mL) than males (0.67 ± 0.13 pg/mL versus 0.75 ± 0.26 pg/mL). IFN-γ concentrations were also lower following Koomba kaat 1 administration (*P* = 0.008), with reductions observed in both males (0.40 ± 0.16 versus 0.86 ± 0.46 pg/mL) and females (0.58 ± 0.39 versus 0.92 ± 0.30 pg/mL). Conversely, IL-12(p40) concentrations were increased (*P* = 0.016), with the greatest increase observed in males (148.93 ± 58.33 pg/mL versus 74.89 ± 24.12 pg/mL). Despite these statistically significant differences, no corresponding changes were detected in BALF cellularity, total macrophage or neutrophil counts, clinical observations, organ pathology or other measures of pulmonary injury.

## Discussion

The successful clinical translation of bacteriophage therapy depends not only on demonstrating antibacterial efficacy but also on establishing the safety of individual therapeutic candidates. Despite increasing clinical use of phages, comprehensive preclinical safety data remain limited, particularly for respiratory administration. Using complementary human airway epithelial and murine models, we demonstrated that high-purity Koomba kaat 1 and Biyabeda mokiny 1 were well tolerated following high-dose respiratory exposure, producing no evidence of epithelial injury, clinically meaningful pulmonary inflammation, or systemic toxicity. Together, these findings provide important preclinical evidence supporting the continued development of these phages for respiratory applications.

A major strength of this study is the use of differentiated primary airway epithelial cells cultured at air-liquid interface (ALI), which more closely replicate the structure and function of the human airway than conventional submerged monolayer cultures (Martinovich et al., 2017; Pezzulo et al., 2011; Trend et al., 2018). Unlike immortalised cell lines, ALI cultures maintain epithelial polarity, mucus production and barrier function, enabling physiologically relevant assessment of host responses to inhaled therapeutics (Silva et al., 2023; Skronska-Wasek et al., 2021). Within this model, neither Koomba kaat 1 nor Biyabeda mokiny 1 altered epithelial morphology, barrier integrity, mucus production, cytotoxicity or IL-8 secretion following high-dose exposure. Although Koomba kaat 1 induced a modest increase in apical IL-8 compared with its bacterial comparator, this was not accompanied by epithelial injury, increased LDH release or changes in basolateral cytokine secretion, suggesting the response was unlikely to be biologically significant.

Mucus production is a critical component of airway defence and represents an underexplored endpoint in respiratory phage research. Previous studies have demonstrated that bacteriophages can adhere to mucosal surfaces and contribute to antimicrobial defence through the bacteriophage adherence to mucus (BAM) model (Barr et al., 2013; Chin et al., 2022). The absence of changes in mucus production following exposure to either phage indicates that purified preparations do not stimulate mucus hypersecretion or disrupt epithelial secretory function, further supporting their compatibility with the respiratory mucosa.

These *in vitro* findings were reinforced by repeated respiratory administration *in vivo*. Mice tolerated twice-daily intranasal administration for 14 days without adverse clinical signs, behavioural changes or weight loss. Food and water consumption were unaffected, and comprehensive necropsy identified no treatment-related abnormalities or changes in organ weights. Although the liver is recognised as a major site of viral clearance (Huh et al., 2019; Sánchez Romano et al., 2024), repeated respiratory exposure produced no evidence of hepatic or systemic toxicity. These observations are consistent with previous studies reporting favourable safety profiles for therapeutic phages (Huang et al., 2022; Lehman et al., 2019) while extending these findings specifically to repeated respiratory delivery of purified *Staphylococcus aureus* phages. Similarly, blood gas, biochemical and haematological analyses demonstrated few treatment-associated effects. The reduction in blood glucose observed in males receiving Biyabeda mokiny 1 represented the only significant biochemical difference but remained within the expected physiological range for C57BL/6 mice (Han et al., 2008) and occurred without corresponding changes in clinical observations, pathology or inflammatory markers. Together, these findings indicate that repeated intranasal administration of either phage does not induce clinically meaningful systemic toxicity.

Assessment of BALF further demonstrated that repeated respiratory administration did not induce pulmonary inflammation. Neither phage altered total BALF cellularity or macrophage and neutrophil numbers, while eosinophils and other inflammatory cell populations were absent across all treatment groups. BALF cellularity is a sensitive indicator of pulmonary inflammation and is widely used to characterise responses to inhaled toxicants, respiratory pathogens and experimental therapeutics (Eisfeld et al., 2019; Fernandes de Souza et al., 2023; Huaux et al., 2002; Landwehr et al., 2023).

Although differential BALF cell counts are commonly reported in respiratory toxicology, they have rarely been incorporated into preclinical evaluations of therapeutic *Staphylococcus aureus* phages. Their inclusion in the present study therefore provides additional evidence that repeated exposure to purified phage preparations does not provoke measurable pulmonary inflammation. BALF protein concentrations likewise supported preservation of epithelial integrity. Although Biyabeda mokiny 1 produced a modest reduction in total BALF protein, this occurred in the absence of epithelial injury, inflammatory cell infiltration or histopathological abnormalities. As elevated BALF protein is a recognised marker of epithelial barrier disruption and vascular leakage, the absence of increased protein concentrations is consistent with a lack of pulmonary toxicity.

Comprehensive cytokine profiling also demonstrated a favourable immunological safety profile. Neither phage altered the principal pro-inflammatory mediators IL-1α, IL-6, G-CSF, KC or TNF-α. While Koomba kaat 1 produced statistically significant changes in IL-17, IFN-γ and IL-12(p40), these differences remained small, occurred at very low absolute concentrations and were not accompanied by changes in BALF cellularity, epithelial integrity or clinical outcomes. Interpretation of multiplex cytokine assays requires consideration of biological as well as statistical significance, as highly sensitive assays frequently detect small differences without physiological consequence (Gueders et al., 2009). For example, IFN-γ concentrations remained below 1 pg/mL, substantially lower than levels associated with inflammatory activation (Wu et al., 2001), while IL-12(p40) concentrations were considerably lower than those reported during experimental pulmonary inflammation or active *S. aureus* infection (Huaux et al., 2002; Peignier et al., 2024). Although IL-12(p40) functions as a chemoattractant for macrophages and activated dendritic cells (Cooper & Khader, 2007), no corresponding increase in macrophage recruitment was observed. Collectively, these findings indicate that the observed cytokine differences represent minor biological variation rather than adverse host responses.

The overall safety profile observed here is consistent with the growing body of evidence supporting the tolerability of therapeutic bacteriophages (Lehman et al., 2019; Liu et al., 2021). Previous studies have primarily focused on efficacy in infection models, including pneumonia, infective endocarditis and septicaemia, where phages have generally been well tolerated while improving bacterial clearance (Prazak et al., 2022; Save et al., 2022; Takemura-Uchiyama et al., 2014; Valente et al., 2021). In contrast, relatively few investigations have evaluated the intrinsic effects of purified phages in the absence of infection or incorporated comprehensive respiratory safety endpoints. By integrating differentiated human airway epithelial cultures with repeated-dose respiratory administration in vivo, this study provides one of the most comprehensive evaluations of host responses to inhaled therapeutic phages reported to date.

Several limitations should be acknowledged. First, safety was assessed in healthy airway models without concurrent bacterial infection; host responses may differ in the presence of active infection owing to interactions between phages, bacteria and the immune system. Second, although the 14-day treatment regimen reflects clinically relevant therapeutic durations (Petrovic Fabijan et al., 2020), future studies should evaluate longer-term dosing, pharmacokinetics and adaptive immune responses. Despite these limitations, this study addresses an important gap in the phage therapy literature by establishing a comprehensive framework for preclinical respiratory safety assessment. The integration of differentiated primary airway epithelial cultures with repeated-dose *in vivo* exposure, pulmonary physiology, BALF cellularity and multiplex cytokine profiling provides a robust platform for evaluating candidate therapeutic phages before clinical application. As inhaled phage therapy advances towards routine clinical use, incorporation of standardised respiratory safety endpoints alongside genomic and functional characterisation will improve the consistency, comparability and translational value of future preclinical studies.

In conclusion, high-purity Koomba kaat 1 and Biyabeda mokiny 1 were well tolerated following repeated respiratory administration and produced no evidence of clinically meaningful epithelial injury, pulmonary inflammation or systemic toxicity. These findings support the continued clinical development of both phages for the treatment of *Staphylococcus aureus* respiratory infections and demonstrate the importance of comprehensive respiratory safety assessment as a critical component of preclinical phage therapeutic development. More broadly, this study provides a practical framework for evaluating the respiratory safety of candidate therapeutic bacteriophages, supporting the translation of inhaled phage therapy into early-phase clinical trials.

## Acknowledgments

Member institutions of Phage WA include the Wal-yan Respiratory Research Centre, Kids Research Institute Australia, The University of Western Australia, Perth, Western Australia, Australia, and St John of God Hospital, Subiaco, Western Australia, Australia. This study was supported by a MRFF Grant (2023559) as well as funding from the Stan Perron Charitable Foundation, the Conquer CF Foundation, and a Department of Health WACRF grant. A.K. is a CFWA/Stan Perron Fellow and S.M.S. was an NHMRC Practitioner Fellow holding an NHMRC Investigator Grant (2007725). We also acknowledge that this project was conducted on the traditional homelands of the Noongar people, with phages isolated from waters across Noongar Wadjak. Phage WA thanks the Sharon Gregory and Walyalap Waangkan Noongar language team, who named the phages in this study in Wadjak Noongar language. Koomba-kaat kep-wari Wadjak (Koomba kaat) translates as “big head-like (from) still water pond Wadjak” and Biyabeda-mokiny kep-wari Wadjak (Biyabeda mokiny) translates as “squidlike (from) still water pond Wadjak”. Wastewater samples for phage isolation were provided by the Water Corporation of Western Australia from their Subiaco Wastewater Treatment Plant.

## Supplementary data

**Supplementary table 1:** Haematology, blood gas and blood chemistry parameters across experimental groups separated by sex. The only significant effects seen were higher base excess in males (when compared to females) and lower blood glucose in males treated with Biyabeda mokiny 1 (when compared to sex matched controls). Data shown are mean (± SD) values (n=6 per treatment group), except in the Koomba kaat 1 (Females) group (n=5), where one animal was excluded due to an iStat Alinity sample failure.

| Parameters | PBS control (Males) | Koomba kaat 1 (Males) | Biyabeda mokiny 1 (Males) | PBS control (Females) | Koomba kaat 1 (Females) | Biyabeda mokiny 1 (Females) |
| --- | --- | --- | --- | --- | --- | --- |
| Na (mmol/L) | 144.67 (± 2.34) | 146.00 (± 3.35) | 147.00 (± 1.90) | 146.33 (± 1.63) | 148.00 (± 1.00) | 146.67 (± 2.42) |
| K (mmol/L) | 4.05 (± 1.34) | 3.47 (± 0.73) | 3.80 (± 0.52) | 3.78 (± 0.29) | 3.84 (± 1.06) | 3.60 (± 0.71) |
| iCa (mmol/L) | 1.08 (± 0.08) | 0.97 (± 0.13) | 1.01 (± 0.08) | 0.99 (± 0.06) | 0.97 (± 0.20) | 0.94 (± 0.13) |
| Glucose (mmol/L) | 22.37 (± 2.13) | 20.20 (± 1.16) | **17.98 (± 2.33) \*** | 20.52 (± 2.74) | 18.45 (± 1.89) | 21.97 (± 3.02) |
| Hct (%PCV) | 38.67 (± 2.73) | 40.00 (± 2.10) | 37.17 (± 3.43) | 38.17 (± 1.72) | 38.00 (± 1.87) | 38.50 (± 1.38) |
| Hb (g/L) | 131.50 (± 9.22) | 136.17 (± 7.25) | 126.50 (± 11.81) | 129.83 (± 6.18) | 129.00 (± 6.28) | 131.00 (± 4.94) |
| pH | 7.36 (± 0.06) | 7.39 (± 0.15) | 7.36 (± 0.04) | 7.32 (± 0.06) | 7.35 (± 0.03) | 7.42 (± 0.10) |
| PCO<sub>2</sub> (kPa) | 4.68 (± 0.86) | 4.16 (± 1.62) | 4.73 (± 0.86) | 4.52 (± 1.04) | 4.73 (± 1.43) | 3.57 (± 0.87) |
| PO<sub>2</sub> (kPa) | 5.02 (± 1.70) | 6.35 (± 2.65) | 5.38 (± 1.46) | 5.82 (± 1.40) | 5.88 (± 1.41) | 6.52 (± 4.32) |
| HCO<sub>3</sub> (mmol/L) | 19.52 (± 2.45) | 17.70 (± 3.07) | 19.98 (± 2.45) | 17.53 (± 3.39) | 17.48 (± 2.94) | 16.72 (± 1.42) |
| base excess (mmol/L) | -6.17 (± 2.64) | -7.33 (± 3.20) | -5.50 (± 2.07) | -8.50 (± 3.73) | -8.20 (± 3.03) | -7.83 (± 0.75) |
| sO<sub>2</sub> (%) | 65.33 (± 24.24) | 74.17 (± 22.17) | 70.50 (± 14.64) | 72.83 (± 12.81) | 72.80 (± 16.38) | 73.50 (± 13.26) |
| TCO<sub>2</sub> (mmol/L) | 20.50 (± 2.51) | 18.67 (± 3.39) | 21.00 (± 2.83) | 18.67 (± 3.50) | 18.20 (± 3.11) | 17.33 (± 1.37) |

Significant difference indicated with a “\*” are between that cells treatment group and sex-matched controls (p< 0.05).

**Supplementary table 2:** Bronchoalveolar lavage mediators across experimental groups and controls separated by sex. Overall, the only significant effects seen were with respect to IL-12(p40), IL-17, and IFN-γ for mice treated with Koomba kaat 1. Data shown are mean (± SD) mediator levels in BALf from male and female mice exposed to phages Koomba kaat 1, Biyabeda mokiny 1 phages, or controls (n=6 per treatment group).

| Parameters (pg/mL) | PBS control (Males) | Koomba kaat 1 (Males) | Biyabeda mokiny 1 (Males) | PBS control (Females) | Koomba kaat 1 (Females) | Biyabeda mokiny 1 (Females) |
| --- | --- | --- | --- | --- | --- | --- |
| IL-1α | 1.03 (± 0.26) | 0.76 (± 0.21) | 1.10 (± 0.22) | 0.94 (± 0.20) | 0.87 (± 0.31) | 1.03 (± 0.32) |
| IL-2 | 0.63 (± 0.00) | 0.63 (± 0.00) | 0.63 (± 0.00) | 0.63 (± 0.00) | 0.75 (± 0.29) | 1.08 (± 0.70) |
| IL-4 | 0.17 (± 0.00) | 0.26 (± 0.22) | 0.27 (± 0.24) | 0.17 (± 0.00) | 0.17 (± 0.00) | 0.30 (± 0.23) |
| IL-5 | 0.46 (± 0.00) | 1.39 (± 1.66) | 1.21 (± 1.83) | 0.46 (± 0.00) | 1.29 (± 0.95) | 2.63 (± 2.83) |
| IL-6 | 0.18 (± 0.00) | 0.18 (± 0.00) | 0.24 (± 0.16) | 0.22 (± 0.12) | 0.54 (± 0.64) | 0.28 (± 0.26) |
| IL-10 | 1.73 (± 0.00) | 2.25 (± 1.28) | 1.73 (± 0.00) | 1.73 (± 0.00) | 1.73 (± 0.00) | 1.73 (± 0.00) |
| IL-12(p40) | 74.89 (± 24.12) | **148.93 (± 58.33) \*** | 76.77 (± 13.47) | 96.75 (± 52.62) | 88.63 (± 22.36) | 100.22 (± 42.75) |
| IL-12(p70) | 4.39 (± 2.11) | 3.11 (± 1.31) | 5.04 (± 1.36) | 4.63 (± 1.53) | 3.24 (± 2.18) | 4.25 (± 1.06) |
| IL-13 | 7.39 (± 0.00) | 7.39 (± 0.00) | 8.97 (± 3.87) | 7.39 (± 0.00) | 7.39 (± 0.00) | 8.69 (± 3.19) |
| IL-17 | 0.75 (± 0.26) | **0.67 (± 0.13) \*** | 0.84 (± 0.26) | 0.83 (± 0.11) | **0.52 (± 0.24) \*** | 0.94 (± 0.33) |
| Eotaxin | 4.62 (± 1.49) | 3.00 (± 0.97) | 3.23 (± 1.10) | 4.42 (± 2.79) | 9.91 (± 12.74) | 4.35 (± 2.06) |
| G-CSF | 3.74 (± 1.69) | 3.56 (± 1.25) | 3.57 (± 1.27) | 3.05 (± 0.00) | 7.23 (± 5.53) | 4.24 (± 2.91) |
| IFN-γ | 0.86 (± 0.46) | **0.40 (± 0.16) \*** | 1.09 (± 0.21) | 0.92 (± 0.30) | **0.58 (± 0.39) \*** | 0.85 (± 0.37) |
| KC | 4.00 (± 1.31) | 4.33 (± 1.18) | 4.68 (± 2.41) | 3.52 (± 1.18) | 6.47 (± 4.48) | 4.12 (± 1.51) |
| MIP-1α | 0.31 (± 0.18) | 0.54 (± 0.41) | 0.40 (± 0.24) | 0.41 (± 0.27) | 0.63 (± 0.38) | 0.59 (± 0.42) |
| MIP-1β | 2.09 (± 0.00) | 2.78 (± 1.71) | 4.27 (± 5.36) | 2.96 (± 2.15) | 8.65 (± 10.91) | 5.65 (± 6.08) |
| RANTES | 4.16 (± 1.54) | 2.95 (± 1.38) | 3.98 (± 0.82) | 3.79 (± 0.51) | 3.87 (± 0.52) | 4.18 (± 2.01) |
| TNF-α | 3.09 (± 1.52) | 2.13 (± 0.92) | 2.34 (± 1.43) | 3.01 (± 1.38) | 3.21 (± 2.37) | 4.54 (± 2.48) |

Significant difference indicated with a “\*” are between that cells’ treatment group and sex-matched controls (p< 0.05).

